# Molecular signaling involved in immune system activation against root-knot nematodes by bio-control agents in tomato plants

**DOI:** 10.1101/556175

**Authors:** Sergio Molinari, Paola Leonetti

## Abstract

The expression of key defense genes was detected in roots and leaves of tomato plants until the 12^th^ day after treatments with a mixture of beneficial bio-control agents (BCAs), as soil-drenches. The expression of the same genes was monitored in pretreated plants at the 3^rd^ and 7^th^ day since the inoculation with the root-knot nematode *Meloidogyne incognita.* Genes dependent on SA-signaling, such as the Pathogenesis Related Genes, *PR1, PR3,* and *PR5,* were systemically over-expressed at the earliest stages of BCA-root interaction. BCA pre-treatment primed plants against root-knot nematodes. The expression of *PR-*genes and of the gene encoding for the enzyme 1-aminocyclopropane-1-carboxylic acid (ACC) oxidase (*ACO*), which catalyzes the last step of ethylene biosynthesis, was systemically enhanced after nematode inoculation in primed plants. Defense related enzyme activities, such as endochitinase and glucanase, were higher in roots of BCA-treated than in those of untreated plants, as well. On the contrary, the expression of genes dependent on JA/ET-signaling, such as Jasmonate Ethylene Response Factor 3 (*JERF3*), did not increase after nematode inoculation in primed plants. The antioxidant system, as indicated by catalase gene expression and ascorbate peroxidase activity, was repressed in infected colonized roots. Therefore, Systemic Acquired Resistance (SAR), and not Induced Systemic Resistance (ISR), is proposed as the molecular signaling that is activated by BCA priming at the earliest stages of root-nematode interaction. Such BCA-induced activation of the plant immune system did not directly act against nematode motile juveniles penetrating and moving inside the roots. It resulted in a drastically decreased number of sedentary individuals and, then, in an augmented ability of the plants to contrast feeding site building by invasive juveniles.

## Introduction

Bio-control agents (BCAs) are beneficial soil-borne micro-organisms that interact with roots and improve plant health. These root-associated mutualists can be divided into three main groups: Bio-control Fungi (BCF), Arbuscular Mycorrhizal Fungi (AMF), and Plant Growth Promoting Rhizobacteria (PGPR) [1, 2]. BCF include the well-studied*Trichoderma* spp.,a class of opportunistic fungi that may colonize roots of most plants, reducing the infection of plant pathogens and parasites and promoting positive responses in stressed plants. AMF are obligate root symbionts, diffused in most of the soils, that improve plant growth and can alleviate both abiotic and biotic plant stresses. Several genera of the rhizosphere bacteria, such as *Pseudomonas* spp., *Bacillus* spp., and *Streptomyces* spp., can enhance plant growth and improve health. BCAs can suppress pests and diseases by activation of plant immune system [1, 2, 3, 4, 5, 6].

Immune response in plants is regulated by several low molecular weight molecules known as phytohormones, i.e. salicylic acid (SA), jasmonic acid (JA) and ethylene (ET). Furthermore, phytohormones regulate many aspects of plant life, as well, such as reproduction and seed production, photosynthesis, flowering, and response to environmental abiotic challenges.BCAs adopt severalsophisticated molecular mechanisms to activate plant immune response against pathogen and parasite attacks. One of the most studiedmechanism is recognized as systemic acquired resistance (SAR), which is otherwise triggered by local infections causing tissue necrosis [7]. SAR provides long-term resistance to (hemi)biotrophic pathogens and pests, is correlated with the activation of Pathogenesis Related (*PR-*) genes, and is mediated by SA.

Rhizobacteria-induced systemic resistance (ISR) is regulated by JA and ET, is not associated with changes in *PR-*gene expression, and is mainly effective against necrotrophic pathogens and herbivorous insects [1, 6]. AMF produce a mycorrhiza-induced resistance (MIR), and like SAR, acts through SA-dependent defenses giving protection against (hemi)biotrophic pathogens and parasites [5]. Although some reports have indicated that MIR might be associated with priming of JA-regulated responses [8], the exact contribution of JA-signaling to MIR has yet to be actually proved, and may be determined by ISR-eliciting rhizobacteria in the mycorrhizosphere [5]. BCF-induced plant resistance has been extensively described, although the signaling elicited seems to vary according to the considered beneficial fungus and the elicited plant species [2]. In a recent study on the interaction of two *T. harzianum* strains (T908, T908-5) with tomato plants, SAR-marker gene expression was markedly repressed as soon as 24 h after fungal inoculation; however, subsequent inoculation with root-knot nematodes (RKNs) caused an over-expression of the same genes [9]. Preconditioning of plant tissue to trigger effective defenses, only when challenged by a/biotic factors, is a suitable strategy generally adopted by plants to save the costs of a permanent activated state, a phenomenon known in literature as priming [10]. Accordingly, some *Trichoderma* spp. probably prime plants for SAR, but the entire pathway is maintained unexpressed until a subsequent pathogen/parasite attack occurs. The same events were reported to occur in cucumber primed by *T. asperellum* (T203) against *Pseudomonas syringae* pv. *lachrymans* [11]. Priming for defense seems to be induced also by AMF [8].

RKNs are obligate soil-borne animal parasites of almost all crops world-wide. They cause significant damages to the attacked crops, and the consequent decrease in both yield and quality leads to economic losses estimated in more than €80 billion/year in worldwide agriculture [12]. RKNs enter the roots as motile second-stage juveniles (J2s), and move intercellularly through the elongation zone to reach some few cortical cells which are thus transformed into discrete giant or nurse cells. Throughout their life cycle, nematodes maintain these elaborate feeding sites that principally serve to actively transfer solutes and nutrients to the developing nematode. J2s soon become sedentary and, through two molts as J3 and J4, develop into adult gravid females. Females parthenogenetically reproduce by laying 200-400 eggs in an external gelatinous matrix, that is clearly visible outside the roots as an egg mass. Moreover, nematode action induces hypertrophy and hyperplasia of the surrounding tissues, thus causing the formation of the familiar galls on roots [13]. RKNs produce several proteins in the esophageal glands that are introduced, via the stylet, into root cells, or transferred to the root apoplasm by secretion from cuticlin or amphids. An increasing amount of reports has shown that most of these proteins are effectors that contribute to plant defense suppression during infection [14, 15]. Control of plant parasitic nematodes is generally difficult and, at present, still relies on the use of chemical toxic nematicides on cash crops. Such large use is increasingly being banned by European Union Directives, with the aim to reduce pesticide contamination of soils and food. Therefore, scientists are looking for alternative low-impact methods of nematode control, such as genetic and induced resistance, or the use of biocontrol agents [16, 17, 18].

Many reports have shown that beneficial root endophytes, such as *Trichoderma* spp., can reduce infections of endoparasitic nematodes through elicitation of the plant immune system [9, 19, 20]. AMF have been reported to be effective against many nematode species [21]. Moreover, it has been shown that MIR involves priming of defense gene responses against RKNs [22].

Rhizobacteria belonging to specific strains of *Pseudomonas* spp. have long been known to be effective in reducing RKN infection through elicitation of ISR [23]. More recently, three strains of *Bacillus subtilis* and one of *Rhizobium etli*, antagonists also of fungal pathogens, have been reported to reduce the number of both galls and egg masses in roots of tomato plants inoculated with RKNs by eliciting ISR [24].

A mixture of AMF, BCF and PGPR was used in this study as a pre-treatment of tomato plants before inoculation with *M. incognita.* Genomic and proteomic techniques were applied to have information on the molecular mechanisms involved in the activation of plant immune system against these soil-borne parasites. We monitored the expression of six genes from both leaves and roots: five involved in defense mediated by different hormones (i.e. SA, JA, ET), and one gene encoding for the antioxidant enzyme catalase. Detection of gene expressions were performed at 3, 7, 8, and 12 days after treatment (dpt) and 3-7 days after inoculation (dpi) with nematodes. Furthermore, we tested key enzyme activities of roots involved in biotic challenges. Therefore, we detected the early response of plants to colonization of beneficial microorganisms, and the priming process that such colonization induces against the subsequent RKN attack. Data of this paper confirm that plant defense against RKNs was activated by the used BCAs, basically through the over-expression of the SA-dependent *PR-*genes.

## Materials and Methods

### Treatments of tomato plants with BCAs

Seeds of the tomato (*Solanum lycopersicum* L.) cultivar Roma VF, susceptible to root-knot nematodes (RKNs) were surface-sterilized and sown in river sand (previously sterilized by autoclaving twice at 121 °C for 30 min). Seedlings were transplanted to 110-cm^3^ clay pots, filled with 150 g of sterilized sand river. Pots were put in temperature-controlled benches (soil temperature 23-25°C), located inside a glasshouse. Plantlets were provided with a regular regime of 12 h light/day, periodically watered and weekly fertilized with Hoagland’s solution. Plants were allowed to grow to the 4-6 compound leaf stage. Before treatments, average fresh weights of plants were measured; young plants with a weight ranging 3-4 g were selected. BCAs contained in Micosat F^®^ (named Myco in the text), a commercial product by C.C.S. Aosta, Italy, were provided to plants at the dosage of 0.2 g product per g plant fresh weight (0.6-0.8 g/plant). One gram Myco is constituted by 40% roots hosting arbuscular mycorrhiza forming fungi of *Glomus* spp. (*Glomus* spp. *GB 67, G. mosseae GP11, G. viscosum GC 41*) and 12.4 x 10^7^ C.F.U. of a mixture of antagonistic fungi (*Trichoderma harzianum TH 01, Pochonia chlamydosporia PC 50*), rhizo-bacteria such as *Agrobacterium radiobacter AR 39, Bacillus subtilis BA 41, Streptomyces* spp., and yeasts (*Pichia pastoris PP* 59). Myco powder was dissolved in a peptone-glucose suspension (0.7 g ml^-1^), and incubated in an orbital shaker at 25°C for 3 days in dark. In some experiments, 100 µg ml^-1^ Amphotericin B, a potent antifungal compound, was added to the suspension to exclude the effect on plants of the fungal components of the mixture. Then, groups of plants were soil-drenched with suitable amounts of Myco suspension, whilst control plants were provided with the sole peptone-glucose suspension.

### Inoculation of tomato plants with nematodes

Populations of the root-knot nematode *Meloidogyne incognita* (Kofoid *et* White) Chitwood, collected from field and reared in a glasshouse on susceptible tomato, were used for plant inoculation. Females of such a population were identified as *M. incognita* by electrophoretic esterase and malate dehydrogenase isozyme patterns [25]. Invasive second-stage juveniles (J2s) were obtained by incubation of egg masses in tap water at 27°C; 3-day-old J2s were collected and used for inoculation. Five days after Myco treatment, groups of treated and untreated plants were inoculated with 300 J2/plant, other groups were left not inoculated. Inoculation was carried out by pouring 2-4 ml of J2 stirring suspensions into 2 holes made in the soil around the plants. Detection of nematode infection was performed 3, 7, 21, and 40 dpi. Plants were grown in pots filled with sterilized river sand in the experiments in which harvest was predicted to occur 3 and 7 days after nematode inoculation; conversely, plants were grown in pots filled with a mixture of sterilized loamy soil and sand (1:1, v:v) when harvest was predicted at 21 and 40 dpi.

### Detection of nematode infection

The numbers of motile vermiform individuals (second stage, J2s) and sedentary swollen individuals (third and fourth stages, sedentary juveniles, SJs) that had, respectively, penetrated and established into the roots 3 and 7 dpi were determined under a stereoscope after coloration by the sodium hypochloride-acid fucsin method [26]. In the roots harvested 21 and 40 dpi, only adult reproducing females and egg masses were searched and counted. Extraction of swollen females from roots was carried out by incubation with pectinase and cellulase enzyme mixture at 37° C in an orbital shaker to soften the roots. After a brief homogenization in physiological solution, females were collected on a 90 µm sieve and counted under a stereoscope (x 12 magnification). Egg masses (EMs) were colored by immersing, for at least 1 h in a refrigerator, the roots in a solution (0.1 g L^-1^) of the colorant Eosin Yellow; red-colored EMs were then counted under a stereoscope (x 6 magnification). Samples were arranged from roots of 2 plants; root samples were weighed before extractions or colorations. The numbers of nematode stages were expressed per g root fresh weight. Additionally, shoot and root weights of treated and untreated inoculated plants were measured after harvest.

### RNA extraction andquantitative Real-Time Reverse PCR

Tissues (leaves and roots) from untreated and Myco-treated plants were collected 3, 7, 8, and 12 dpt. Tissues from untreated and Myco-treated plants, inoculated with nematodes, were collected 3 and 7 dpi. Tissue sampleswere weighed and stored at −80°C, if not immediately used for RNA extraction. Plants coming from 2 independent bioassays were used; RNA was extracted from 6 different samples of leaves and roots per treatment, harvested at each dpt and dpi. Tissue samples were separately ground to a fine powder in a porcelain mortar in liquid nitrogen. An aliquot of macerated tissue (100 mg per sample) was used for RNA extraction. Extractions of total RNA were carried out using an RNA-easy Plant Mini Kit (Qiagen, Germany), according to the instructions specified by the manufacturer. RNA quality was verified by electrophoresis runs on 1.0% agarose gel and quantified using a Nano-drop spectrophotometer. QuantiTect Reverse Transcripton Kit (Qiagen, Germany) with random hexamers was used for cDNA synthesis, from 1 μg of total RNA, according to the manufacturer’s instructions. Single 20-μl PCRs included 10 μM each of forward and reverse primers, 1.5 μl cDNA template and 10 μl SYBR^®^ Select Master Mix (Applied Biosystems, Italy). PCR cycling consisted in pre-incubation at 95 °C (10 min); 40 cycles at 95 °C (30 s), at 58 °C (30 s), at 72 °C (30 s), with a final extension step at 72 °C (7 min).

qRT-PCRs were performed in triplicate, using an Applied Biosystems^®^ StepOne™ instrument. Actin was used as the reference gene, since its expression in tomato tissues has been proved not to vary after infestation by nematodes. The GenBank accession used for PR-1 was described as *PR-1b* (*P6*) in [27]. Primers for the analyzed genes are described in Table 1. In order to evaluate the relative expression of the analyzed genes in tissues collected from untreated and Myco-treated plants, 1/ΔC_t_ of each reaction was calculated, being ΔC_t_ = C_t_ (test gene) - C_t_ (reference gene); higher the 1/ΔC_t_ values, higher the expressions of tested genes.

**Table 1.**
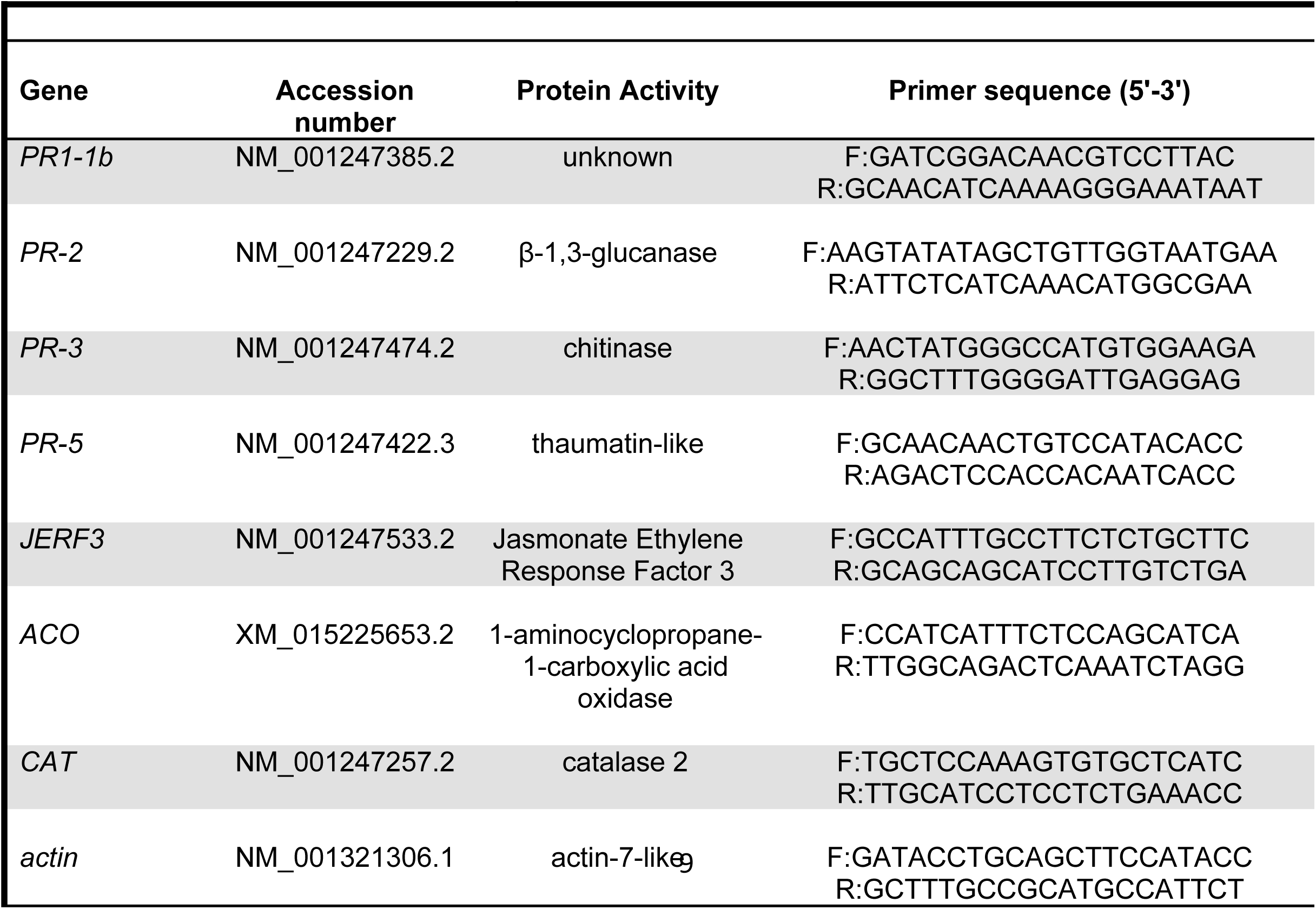
Tomato defense-related genes examined in this study and the specific primers used in quantitative reverse transcriptase-polymerase chain reaction (qRT-PCR)

### Protein extraction and enzyme activity assays

Proteins were extracted from roots of plants at8 and 12 dpt and at 3 and 7 dpi. Roots were set free from sand, and thoroughly rinsed with tap water. Roots and leaves were separated from shoots. Roots from untreated and Myco-treated plants were collected, dried, weighed and put on ice. Root samples were immediately used for protein extractions or stored at −80°C. Samples were ground in porcelain mortars by immersion in liquid nitrogen. For each bioassay, three different powdered samples of roots, coming from 6 plants per treatment, were produced and suspended in a grinding buffer (1:5, w:v) of 0.1 M potassium phosphate buffer (pH 6.0), added with 4% polyvinylpyrrolidone and the protease inhibitor phenyl-methane-sulfonyl fluoride (PMSF, 1 mM). Suspensions were further ground using a Polytron^®^ PT–10–35 (Kinematica GmbH, Switzerland), and filtered through four layers of gauze. Filtrates were centrifuged at 12000 x *g* for 15 min. Supernatants were filtered through 0.45 µm nitrocellulose filters applied to 10-ml syringes. These filtrates were ultra-filtered at 4°C through 20-ml Vivaspin micro-concentrators (10,000 molecular weight cut off, Sartorius Stedim, Biotech GmbH, Germany). Retained protein suspensions were used for protein content and enzyme assays. Protein content was determined by the enhanced alkaline copper protein assay, with bovine serum albumin as the standard [28].

Chitinase activity (CHI) was measured by a colorimetric procedure that detects N-acetyl-D-glucosamine (NAG) [29]. The hydrolytic action of chitinase produces chitobiose which is converted into NAG by the β-glucuronidase introduced in the reaction mixture. Suspended chitin (250 µl, 10 mg/ml) from shrimp shells (Sigma-Aldrich, Italy) was added to 50 µl of leaf extract or 100 µl of root extract diluted in 200-150 µl of 0.05 M sodium acetate buffer (pH 5.2) containing 0.5 M NaCl. The reaction was allowed by incubating the mixtures in eppendorfs for 1 h at 37°C in an orbital incubator, and stopped by boiling at 100°C for 5 min in a water bath. Eppendorfs were centrifuged at 10000 x *g* for 5 min at room temperature. Supernatants (300 µl) were collected and added with 5 µl β-glucuronidase (Sigma, type HP-2S, 9.8 units/ml). Reaction on/off was carried out as previously described; reaction mixtures were let cool at room temperature. After adding 60 µl of 0.8 M potassium tetraborate (pH 9.1), mixtures were heated to 100°C for 3 min and cooled to room temperature. Then, 1% 4-dimethylaminobenzaldehyde (1.2 ml, DMAB, Sigma) was added, and mixtures incubated at 37°C for 20 min. Absorbance was read at 585 nm (DU-70, Bechman), and the amount of NAG produced was determined by means of a standard curve obtained with known concentrations (4.5-90 nmoles) of commercial NAG (Sigma). Blanks (negative controls) were mixtures in which tissue extracts were not added; positive controls were arranged by adding 10 µl chitinase from *Streptomyces griseus* (Sigma, 200 units/g). The assay was conducted on 6 samples per treatment, and chitinase expressed as nanokatal per mg protein (nkat/mg prot), with 1 nkat defined as 1.0 nmol NAG produced per second at 37°C.

β-1,3-Endoglucanase (glucanase, GLU) activity was measured by determining the amount of glucose released from laminarin (Sigma, Italy) used as substrate. Reaction mixtures consisted in laminarin (0.4 mg) and 100 µl tissue extracts in 300 µl 0.1 M sodium acetate (pH 5.2) that was incubated at 37°C for 30 min. After incubation for glucose production, Nelson alkaline copper reagent (300 µl) was added and the mixtures kept at 100°C for 10 min. Once mixtures had cooled at room temperature, Nelson chromogenic reagent (100 µl) was added for reducing sugars assays [30]. Negative and positive controls consisted of grinding buffer and laminarinase (2 U/ml), respectively. Enzyme activity was expressed as µmol glucose equivalents released per minute, according to a standard curve created with known amounts (10-200 µg ml^-1^) of commercial glucose (Sigma, Italy).

Ascorbate peroxidase activity (APX) was determined as the rate of disappearance of ascorbate in presence of hydrogen peroxide [31]. Reaction mixtures contained 0.1 M TES, pH 7.0, 0.1 mM EDTA, 1 mM ascorbate, 0.1 mM H_2_O_2_, 10-20 µl root extracts, in 0.5 ml final volume. Decrease in absorbance at 298 nm was monitored in a double-beam spectrophotometer (PerkinElmer 557) and indicated ascorbate oxidation; 1 unit of enzyme expressed the oxidation of 1 µmole ascorbate per min (ε=0.8 mM^-1^ cm^-1^).

### Statistical analysis

Means of values ± standard deviations of nematode stages found into the roots were calculated by 9 replicates (n=9), coming from 3 different experiments, arranged in 6 plants per treatment. Weight values of roots and shoots are means ± standard deviations from 18 replicates (n=18). Means from untreated and Myco-treated plants were separated by a paired *t-*test (*P<0.05; **P<0.01). As it concerns qRT-PCR data, means ± standard deviations of 1/ΔC_t_ values of each group from untreated and Myco-treated tissues (n=6) were separated by the non-parametric Kolmogorov-Smirnov test (*P<0.05). As it concerns enzyme activity values, means ± standard deviations were the result of 9 replicates (n=9). Nine tissue samples were obtained from 3 different bioassays. Moreover, each value was calculated on the basis of 3 repeated spectroscopic measurements on each protein extract. Values of enzyme activities were expressed as units mg^-1^ protein; means were separated by a paired *t-*test (\**P*<0.05; \*\**P*<0.01).

## Results

### BCAs activate the immune response of tomato plants

Expression of six genes involved in defense to biotic challenges were detected by qRT-PCR in roots and leaves of plants 3, 7, 8, and 12 dpt with Myco, a commercial product containing a mixture of AMF, BCF, and PGPR. At first, 3 genes, *PR1, PR3,* and *PR5*, were tested*. PR1-P6* or *PR1b1* encodes for one of the PR-1 protein subfamily, which consists of low molecular-weight proteins of unknown biochemical function. We chose to test *PR1b1* gene expression because it was found to be strongly activated during the hypersensitive response (HR) to pathogens in tomato, whilst the other gene of the family, *PR1a2*, was not induced by pathogenic signals [32]. *PR3* gene encodes for several types of endochitinases, and has been reported to be induced by ethylene treatments in tomato [33]. *PR5* gene family encodes for thaumatin-like proteins and is involved in osmotic regulation of cells. Expression of *PR1* and *PR5* are highly induced by SA accumulation and over-expressed in SAR against biotrophic pathogens [34]. Expression of *PR1* gene was highly activated in leaves and roots from Myco-treated plants, as soon as 7 dpt. After this early activation, *PR1* expression in treated plants was found to be repressed with respect to untreated plants (Fig. 1A-B). No significant changes in *PR3* gene expression between untreated or treated plants were observed up to 8 dpt; at 12 dpt, a significant inhibition of the gene expression was detected in both roots and leaves due to Myco treatment (Fig. 1C-D). Activation of *PR5* gene expression was delayed to 8-12 days after Myco treatment and occurred only in roots (Fig. 1E); conversely, in leaves, *PR5* gene seems to be down-loaded in the later stages of the experimental period (Fig. 1F).

**Figure 1.**
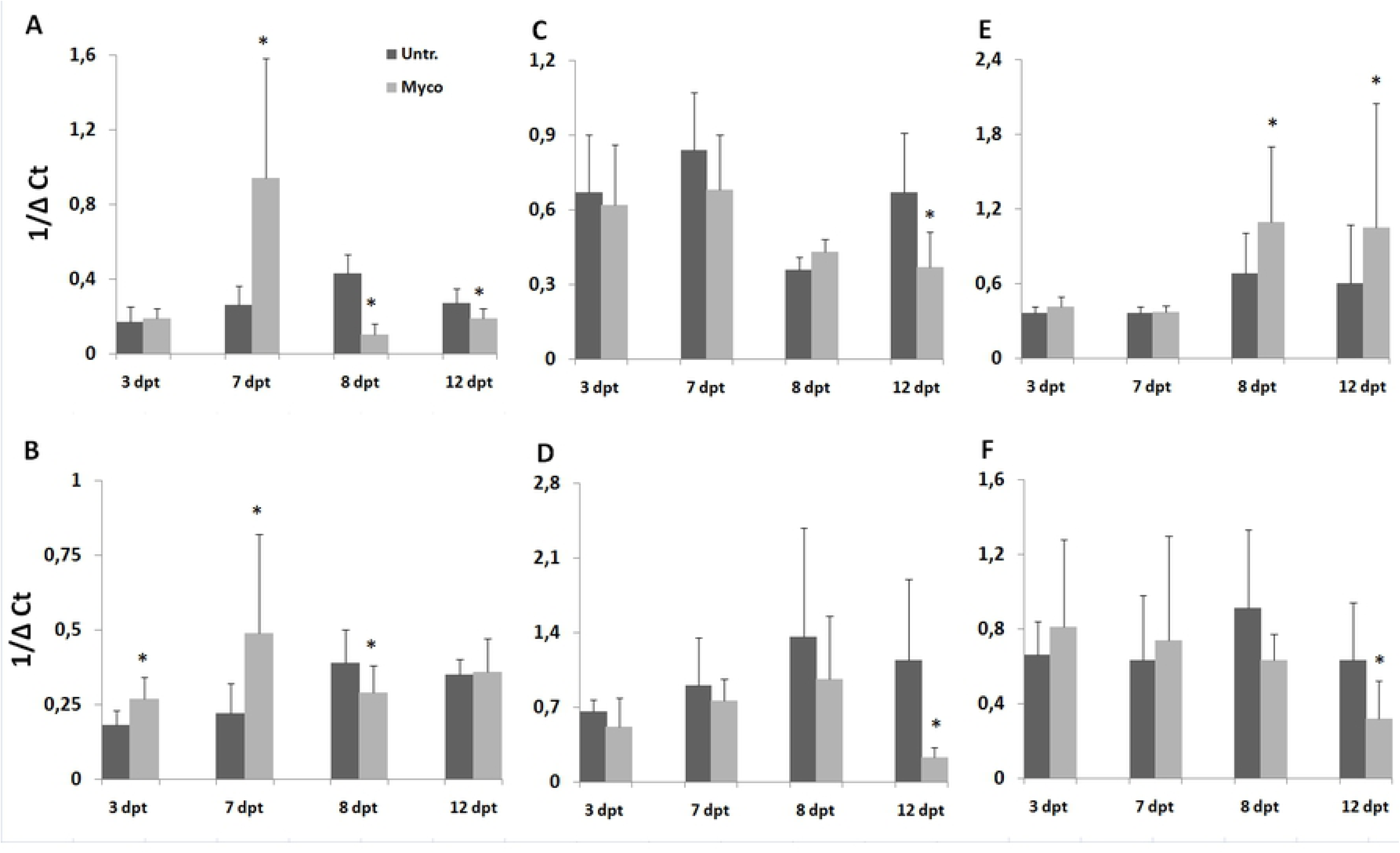
Expression of *PR1, PR3,* and *PR5* genes in tomato tissues after treatment with BCAs. BCAs were provided to tomato plants as Myco soil drenches. Untreated (Untr.) and Myco-treated (Myco) plants are compared. qRT-PCRs were performed to determine ΔC_t_ of *PR1, PR3, PR5* genes in roots (A, C, E, respectively) and leaves (B, D, F, respectively). Tissues were collected 3, 7, 8, 12 days after Myco treatments (dpt). Values are expressed as 1/ΔC_t_ means ± standard deviations. Means are separated by the non-parametric Kolmogorov-Smirnov test (*P<0.05).

The second series of 3 genes tested included Jasmonate Ethylene Response Factor 3 (*JERF3*), the gene encoding for the enzyme 1-aminocyclopropane-1-carboxylic acid (ACC) oxidase (*ACO*), and the gene encoding for the enzyme catalase (*CAT*). *JERF3* encodes for a member of ERF proteins, a trans-acting factor responding to both ET and JA in tomato [35]. ACC oxidase is the enzyme which catalyzes the last step of ET biosynthesis, whilst catalase is one of the key enzyme of the antioxidant enzyme system which neutralizes the toxic hydrogen peroxides produced in plant defense against pathogens and parasites. *JERF3* gene is significantly downloaded in Myco-treated plants at 8 and 12 dpt (Fig. 2A-B). Expression of *ACO* gene is not generally affected by treatment with Myco; however, its expression in tomato plants consistently decreased after 7 dpt (Fig. 2C-D). This reduction in expression at later times occurred also for *CAT* gene; however, Myco-treated plants showed an over-expression of *CAT* gene at earlier times after treatment (3-7 dpt, Fig. 2E-F).

**Figure 2.**
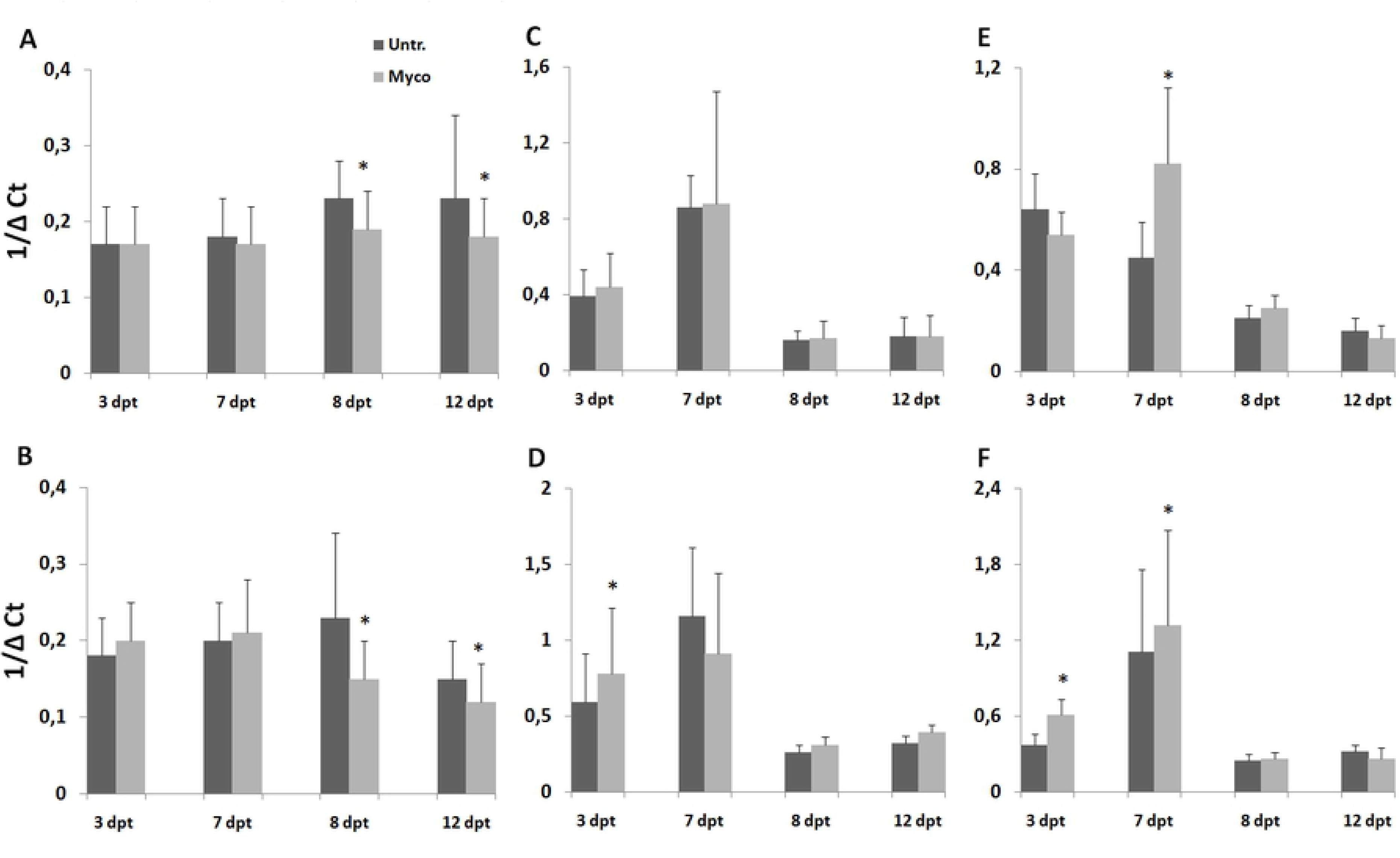
Expression of *JERF3, ACO,* and *CAT* genes in tomato tissues after treatment with BCAs. BCAs were provided to tomato plants as Myco soil drenches. Untreated (Untr.) and Myco-treated (Myco) plants are compared. qRT-PCRs were performed to determine ΔC_t_ of *JERF3, ACO*, and *CAT* in roots (A, C, E, respectively) and leaves (B, D, F, respectively). Tissues were collected 3, 7, 8, 12 days after Myco treatments (dpt). Values are expressed as 1/ΔC_t_ means ± standard deviations. Means are separated by the non-parametric Kolmogorov-Smirnov test (*P<0.05).

### BCAs prime tomato plants against root-knot nematodes

The amount of motile invasive J2 into the roots at 3 and 7 dpi was not significantly affected by BCA treatment. However, feeding site construction is the early step of infection, at which motile J2 become sedentary and start to grow and transform cortical cells into nursery cells, that transfer nutrients from plant metabolism to the developing nematodes. At 7 dpi, sedentary juveniles extracted from roots of Myco-treated plants were one third of those from untreated plants. At 21 dpi, Myco treatment caused a high decrease of the numbers of reproducing females and egg masses present in/on roots. At the end of life cycle of successfully developed nematodes (40 dpi), females and egg masses in roots of Myco-treated plants were still significantly lower than in roots of untreated plants, although at a minor extent. When Myco suspensions were added with the potent antifungal compound Amphotericin B, the suppressive effect of the BCA mixture on nematode infection was inverted; inactivation of the fungal components resulted in a significant augment of females and egg masses in Myco-treated with respect to untreated plants (Table 2).

**Table 2.** Nematode individuals penetrated, developed and reproduced in roots of tomato untreated and treated with Myco at different days after inoculation (dpi)

| dpi | average no. per plant ± stdev | | | | average no. per g root fresh weight | | | | | | | |
| --- | --- | --- | --- | --- | --- | --- | --- | --- | --- | --- | --- | --- |
|  | Shoot Weight (g) |  | Root Weight (g) |  | Motile invasive J2 |  | Sedentary J3-4 forms |  | Females |  | Egg masses |  |
|  | Untreated | Treated | Untreated | Treated | Untreated | Treated | Untreated | Treated | Untreated | Treated | Untreated | Treated |
| 3 | 2.5±0.6 | 2.4±0.5 | 0.4±0.2 | 0.4±0.2 | 18±10 | 14±10 | 0 | 0 | 0 | 0 | 0 | 0 |
| 7 | 3.3±0.8 | 3.3±0.6 | 0.4±0.2 | 0.4±0.2 | 146±74 | 112±84 | 24±12 | 8±7* | 0 | 0 | 0 | 0 |
| 21 | 4.8±1.7 | 4.9±1.3 | 1.2±0.6 | 1.1±0.8 | nd<sup>b</sup> | nd | nd | nd | 28±12 | 6±4* | 12±5 | 2±2* |
| 40 | 9.4±3.6 | 8.9±3.8 | 1.8±1.0 | 2.2±1.3* | nd | nd | nd | nd | 155±28 | 83±10* | 97±28 | 52±14* |
| 40+AMPHO<sup>a</sup> | 11.4±2.5 | 11.7±2.5 | 1.8±0.8 | 1.9±0.8 | nd | nd | nd | nd | 169±67 | 378±155* | 102±33 | 168±18* |

\* significantly different ( $P < 0.05$ ) according to a paired $t$ -test; <sup>a</sup>tests in which Myco suspension was added with 100 $\mu\text{g ml}^{-1}$ Amphotericin B; nd=not determined

The occurrence in tomato plants of the priming phenomenon, induced by the BCAs used in this study as a pre-treatment, is indicated by the over-expression of *PR-*genes at 3 and 7 days after nematode inoculation of pre-treated plants. Gene over-expression involved all the tested *PR-*genes (*PR1, PR3,* and *PR5*), and was systemic (Fig. 3). The only exception was detected for *PR3* gene expression in leaves at 7 dpi (Fig. 3D).

**Figure 3.**
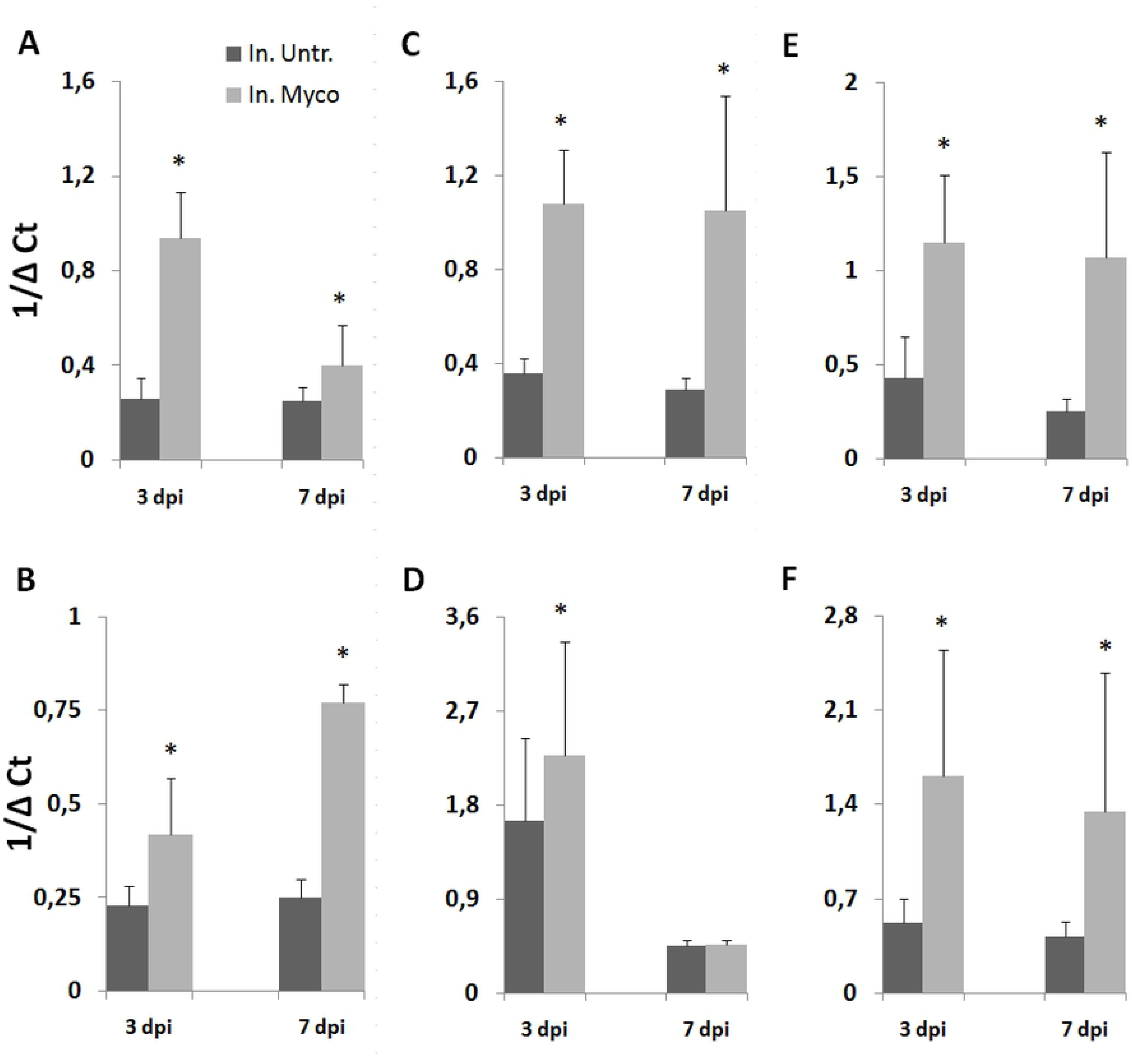
Expression of *PR1, PR3,* and *PR5* genes in tomato tissues of BCA-pretreated plants after inoculation with RKNs. BCAs were provided to tomato plants as Myco soil drenches 5 days before nematode inoculation. Inoculated untreated (In. Untr.) and inoculated Myco-treated (In. Myco) plants are compared. qRT-PCRs were performed to determine ΔC_t_ of *PR1, PR3, PR5* in roots (A, C, E, respectively) and leaves (B, D, F, respectively). Tissues were collected 3 and 7 days after inoculation (dpi). Values are expressed as 1/ΔC_t_ means ± standard deviations. Means are separated by the non-parametric Kolmogorov-Smirnov test (*P<0.05).

Conversely, *JERF3* gene expression of inoculated plants was not affected when plants were treated with Myco, except for a slight but significant decrease, occurring in leaves at 3 dpi (Fig. 4A-B). Slight up-loading of *ACO* gene occurred in Myco-treated plants in roots at 7 dpi and in leaves at both 3 and 7 dpi (Fig. 4C-D). Moreover, *CAT* gene expression was consistently inhibited in roots of inoculated plants by BCAs (Fig. 4E-F).

**Figure 4.**
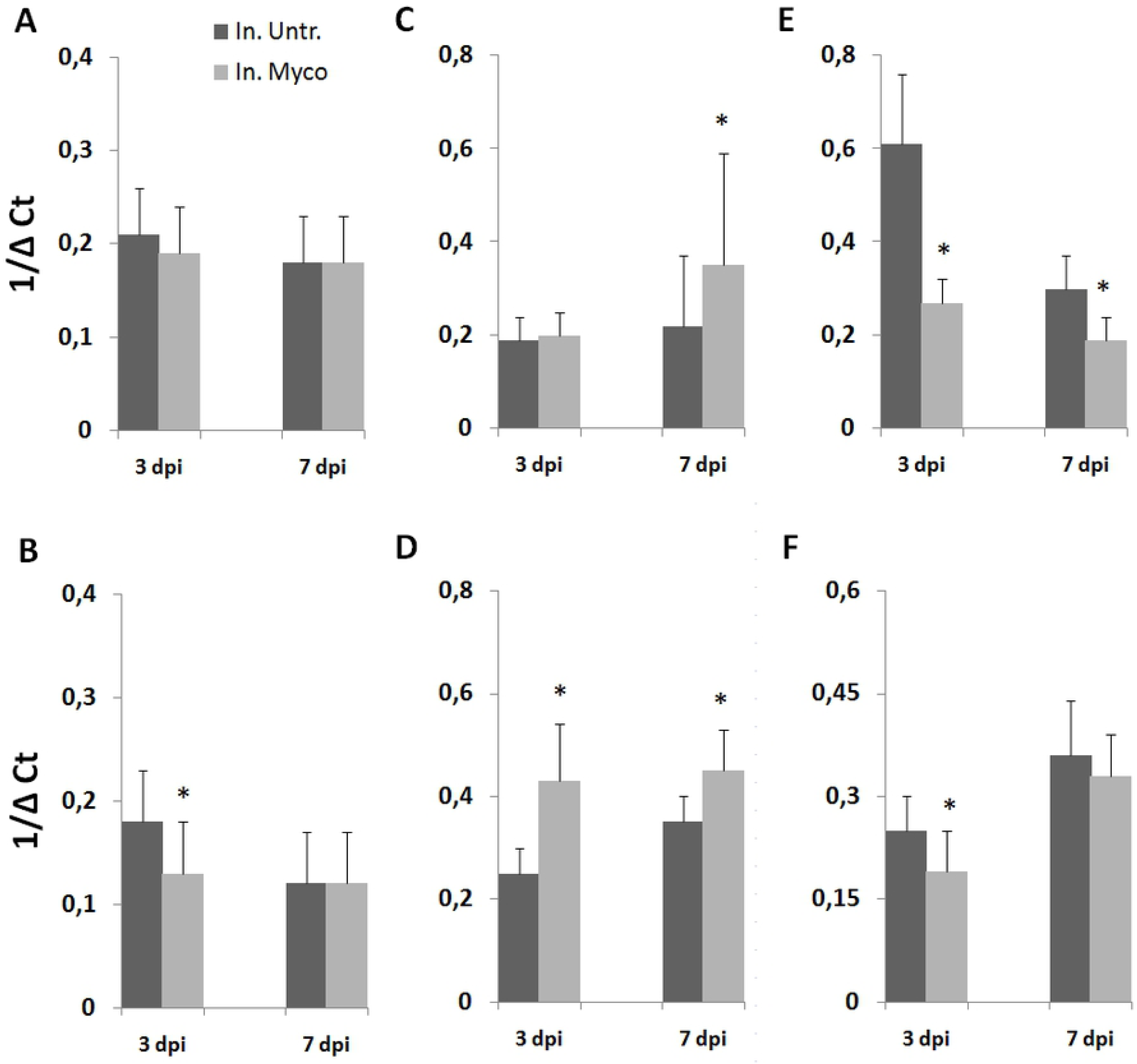
Expression of *JERF3, ACO,* and *CAT* genes in tomato tissues of BCA-pretreated plants after inoculation with RKNs. BCAs were provided to tomato plants as Myco soil drenches 5 days before nematode inoculation. Inoculated untreated (In. Untr.) and inoculated Myco-treated (In. Myco) plants are compared. qRT-PCRs were performed to determine ΔC_t_ of *JERF3, ACO*, and *CAT* in roots (A, C, E, respectively) and leaves (B, D, F, respectively). Tissues were collected 3 and 7 days after inoculation. Values are expressed as 1/ΔC_t_ means ± standard deviations. Means are separated by the non-parametric Kolmogorov-Smirnov test (*P<0.05).

Chitinase (CHI) and glucanase (GLU) are defense-induced enzymes in plants. Moreover, Reactive Oxygen Species (ROS), such as hydrogen peroxide (H_2_O_2_), are normally produced in response to biotic challenges because anti-microbial. H_2_O_2_ is presumed to orchestrate basal and systemic defense to invading pests. Antioxidant enzymes, such as ascorbate peroxidase (APX), degrade H_2_O_2_ favoring biotic infections. We tested the activity of these three enzymes in roots of untreated and Myco-treated tomato plants at 3 and 7 dpi (Table 3). CHI activity was moderately induced by nematode infection at both 3 and 7 dpi. When plants were pre-treated with BCAs, a more intense induction of this activity was observed. Conversely, GLU activity seems not to be activated by nematode infection; however, if plants were pre-treated with BCAs, a marked increase (+62%) of this activity was apparent due to nematode infection at 7 dpi. Nematode infection favored the increase of APX activity to maintain low peroxidative reactions which can jeopardize J2 development. Myco pre-treatment was not able to restrain this increment, at least during the earliest stages of infection.

**Table 3.** Effect of RKN inoculation on enzyme activities in roots of tomato plants untreated or pre-treated with Myco at different days after inoculation (dpi)

| Enzyme | Roots untreated | Inoculated Roots untreated | effect % | Roots Myco-treated | Inoculated Roots Myco-treated | effect % |
| --- | --- | --- | --- | --- | --- | --- |
| **CHI<sup>a</sup>** |  |  |  |  |  |  |
| 3 dpi | 0.15±0.02 | 0.19±0.04** | +30 | 0.22±0.07 | 0.32±0.15* | +48 |
| 7 dpi | 0.32±0.11 | 0.37±0.09* | +15 | 0.19±0.06 | 0.28±0.12** | +50 |
| **GLU<sup>b</sup>** |  |  |  |  |  |  |
| 3 dpi | 46.2±8.0 | 48.6±11.1 | ns | 51.8±6.4 | 54.0±11.9 | ns |
| 7 dpi | 51.8±14.5 | 65.5±10.2 | ns | 33.5±10.1 | 54.4±5.9* | +62 |
| **APX<sup>c</sup>** |  |  |  |  |  |  |
| 3 dpi | 0.24±0.03 | 0.40±0.04** | +65 | 0.32±0.03 | 0.47±0.14* | +46 |
| 7 dpi | 0.41±0.16 | 0.54±0.12** | +33 | 0.24±0.03 | 0.37±0.04** | +53 |

significantly different (\*P<0.05; \*\*P<0.01) according to a paired *t*-test; ns= not significant; <sup>a</sup>chitinase expressed as nkat mg<sup>-1</sup> prot; <sup>b</sup> glucanase expressed as μmol glucose min<sup>-1</sup> mg<sup>-1</sup> prot; <sup>c</sup>ascorbate peroxidase expressed as μmole ascorbate min<sup>-1</sup> mg<sup>-1</sup> prot

## Discussion

Most of the BCAs used in this study to induce plant immune system were AMF and BCF. Symbiotic fungi colonize plant roots of dicots and monocots, and such interaction has as a consequence the reprogramming of plant transcriptome and proteome [2]. One of the main effect of changes in gene expression of colonized plants is the elicitation of resistance to a large variety of pathogens and parasites, from fungi to viruses, nematodes included. We analyzed changes in expression of genes involved in plant defense up to 12 days after a soil-drench treatment of tomato plants with both AMF and BCF. Most of the analyzed genes resulted either up- or down-loaded, and the response was generally systemic. *PR1* and *PR5,* markers of SAR induction mediated by SA, were over-expressed after treatment with the BCA mixture, although at different experimental times. It was possible to observe, at least for *PR1* gene, that the initial activation of expression was followed by a drastic inhibition. A systemic repression of gene expression in later stages of BCA-plant interaction involved *PR3* and *JERF3*, as well, although it was not preceded by an activation. Successful colonization seems to rely on an induced inhibition in plants of CHIgene transcripts and enzyme activity, as well as of JA/ET signaling. A higher amount of *CAT* gene transcripts was initially found in BCA-treated than in untreated plants. A putative increase of the H_2_O_2_-neutralizing enzyme catalase might be induced by colonizing BCAs to protect themselves from H_2_O_2_, that plants generally produce in the early response to biotic challenges. However, at later stages of interaction, *CAT* gene expression of plants dramatically decreased and became similar in both untreated and BCA-treated plants. At 12 dpt, APX activity was found to be consistently inhibited in colonized roots, thus suggesting that the antioxidant system may progressively be repressed along with the stabilization of colonization.

The transient nature of defense gene activation during the early stages of BCA-plant interaction is similar to that recorded during the early stages of mycorrhization. It is likely that AMF secrete suppressors of immunity as a strategy which they share with pathogenic fungi. Specifically, AMF repress SA-dependent defense in later stages to achieve a compatible interaction [8]. Comparably, down-loading of SA- and JA-dependent genes was observed in later stages (8 and 12 dpt) of BCA-plant interaction, and was systemic. Conversely, pre-treatment of tomato plants with two selected strains of *T. harzianum* caused a repression of defense gene expression as early as one day after conidia inoculation, that lasted until 15 dpt [9]. In this case, defense gene activation, that should characterize the initial phase of plant reaction to symbiont fungi invasion, may have passed unobserved, as it is known that gene activation may occur as early as only one hour after *Trichoderma* inoculation [2]. In the present study, the colonization process seems much slower, and the time course of gene expression changes has strict similarities with root mycorrhization. Evidently, a continuous monitoring of genome changes induced by beneficial symbionts over time is mandatory to have a complete information about which molecular signaling is involved in root colonization. Both AMF and BCF act like biotrophs on plants, and share similarities with biotrophic pathogens, such as their sensitivity to SA-regulated defenses [36]. SA-dependent signaling has been shown to be important in the reaction of tomato plants to the symbiont microorganisms used in this study, as it generally occurs against biotrophic pathogens [37]. After gene up-regulation by plants to contrast their diffusion, invading beneficial fungi are able to mediate a counteraction and repress defense gene expression. In our system, both SA- and JA/ET-dependent signaling seem to be repressed. However, if plants were subsequently inoculated with RKNs, the expression of defense genes was much higher than if the nematodes or BCAs were used singly, thus proving that BCA pre-treatment primed tomato plants for enhanced defense against RKNs. In the primed state, the immune system of plants is activated, and plants respond to biotic attacks with faster and stronger defense activation [10]. Gene over-expression involved all the *PR-*genes analyzed and *ACO* gene and was observed 3 and 7 dpi, both in roots and leaves. Conversely, the JA-dependent *JERF3* gene was not activated against RKNs, whilst *CAT* gene was repressed.

A more effective defense induced by BCAs against RKNs is substantiated by the lower numbers of every sedentary forms (J3-4, reproducing females) found during the whole infective process in treated plants, compared with controls. Conversely, the amount of migratory invading J2s, which penetrated into the roots at 3 and 7 dpi, did not decrease because of BCA treatment. Evidently, activation of immunity in this type of plant-pest interaction acts by opposing the attempt by the invading J2 to build a feeding site at the expense of few cortical cells in the root elongation zone. A functional feeding site allows the juvenile to suck nutrients from plant metabolism, to become sedentary and develop into a reproducing female. It is now generally recognized that RKNs are able to suppress plant immune system through injection of an array of effectors directly into the cells and/or by secretion from cuticlin or amphids in the root apoplasm [14, 15, 38]. It is evident that the release of these effectors triggers successful defense reactions in primed plants. Elicitation of plant defense machinery in primed plants, in terms of defense gene over-expression, occurred in this study as early as 3 days after inoculation, when only motile forms were found. This study clearly indicates that primed plants perceive nematode effectors and activate their defense to nematode infection already before the arrangement of the first feeding sites. In other words, activated plants start to recognize and respond to parasitic attack when juveniles are still moving through the elongation zone in search of suitable cortical cells to pierce with their stylet for nutrition. According to our findings, plant immunity may be as rapid as to be triggered by contact with nematodes. However, the effect of immunity did not result in a decrease of nematode root penetration, but in a restriction of the number of nematodes able to build their feeding sites and become sedentary. Once the feeding site is successfully arranged by the juvenile, development and reproduction are no longer hampered.

When BCAs were incubated with Amphotericin B, a potent antifungal compound, pre-treatments of plants lost their ability to induce resistance. On the contrary, nematode infection appeared more severe on pre-treated plants, as indicated by a marked increase of adult females and egg masses. It was ascertained that the rhizobacteria present in the mixture were responsible of this reversed effect on nematode infection. Addition of antibiotics in the antifungal-treated mixture annulled the positive effect on nematode infection; the involvement of abiotic factors was ruled out by pre-treating plants with sterilized mixture that did not cause any changes in nematode infection (results not shown). *Agrobacterium radiobacter AR 39, Bacillus subtilis BA 41, Streptomyces* spp. were the rhizobacteria present in the BCAs mixture used in this study. Actually, different strains of *B. subtilis* were recently proven to induce systemic resistance of tomato plants to RKNs [24]. However, it is generally known that strain specificity is crucial for generating ISR. The rhizobacterial strains used in this study were apparently able, in the absence of functional AMF/BCF, to induce susceptibility to RKNs, as it has been described for one strain of *Pseudomonas fluorescens* (WCS417r) tested on *Arabidopsis thaliana* against aphids [39].

Suppression of immune plant system by RKNs is mediated by an extensive down-regulation of gene expression, particularly *PR-*genes [40, 41]. *PR-*gene down-regulation by RKNs in susceptible tomato plants is generally confirmed by our present and previous data [42]. In contrast, BCA priming enables plants to up-regulate *PR-*genes in response to nematode attack. Such up-regulation is comparable to that of corresponding genes in leaves, thus suggesting that there must be a diffusible signal moving from the roots up to the leaves. It can be presumed that the up-regulation of defense genes observed in leaves may be a marker of induced resistance also to aboveground pests and pathogens. Actually, Myco-treated tomato plants have been found to be poorer hosts of the miner insect *Tuta absoluta* with respect to untreated plants (results not published). Considering that most of the over-expressed genes by BCA priming against RKNs are SA-dependent, the described defense mechanism is likely to be assigned to SAR, which is effective against biotrophs [7].

*ACO* gene encodes for the enzyme involved in the last step of ET biosynthesis. In primed plants, nematode infection induces a systemic enhanced expression of this gene, with a predicted increase of ET level in roots and leaves, which might contribute to limit insect and nematode infections. Actually, ET and ET-signaling have already been reported to play a role in plant defense against endoparasitic sedentary nematodes [43]. BCA-induced SA- and ET-signaling may cooperate for a more efficient and rapid response to nematode infection, as recently reported for *Trichoderma-*induced priming [9]. On the other hand, synergistic signaling cross-talks in plant resistance are commonly reported in literature [44]. *JERF3* gene encodes for a nuclear DNA-binding protein which acts as a transcription factor inducing the expression of JA and ET-dependent defense genes [35]. BCA-mediated priming of tomato plants does not seem to involve the activation of this gene. If we consider *JERF3* as a marker gene for the rhizobacteria-mediated ISR, we can reasonably argue that ISR is not activated in primed tomato plants against RKNs. Conversely, JA-mediated ISR is generally known to activate defense against necrotrophs or herbivorous insects [8].

Compatible plant-parasite interaction are characterized by an increased activity of antioxidant enzymes, such as catalase, to maintain low the level in cells of toxic ROS. Expression of *CAT* gene in Myco-treated plants was generally inhibited after nematode inoculation compared with that in untreated plants. A SAR-mediated defense requires SA accumulation in plant cells which induces H_2_O_2_ accumulation [45]. In primed plants, the observed early down-loading of *CAT* gene may lead, in later stages of biotic challenges, to a lower cell activity of catalase, and, consequently, to the maintenance of elevated amount of H_2_O_2_ in challenged tissues. It is possible that, until this inflammatory-like state is maintained, nematode settling inside the roots may be strongly contrasted. For instance, adult females extracted from primed roots 21 days after inoculation were about 80% less than those from not primed control roots. At 40 days after inoculation, much more individuals were found to have developed up to gravid females, also in primed plants. It can be argued that priming can lose its effectiveness over time, and thus, the many living motile juveniles, which had previously entered the roots, may subsequently have the chance to build their feeding site and develop. However, the overall protective effect of priming determines about 50% inhibition of infection at the end of experimental time, in terms of less females and egg masses found in roots.

In conclusion, data presented herein provide evidence that the mechanisms involved in the activation of plant immune system [3] by beneficial fungi against soil-borne parasite, such as RKNs, rely mainly on SA-mediated signaling and SAR. The immunity conferred is systemic but probably limited in time, at least when it is exerted against RKNs. Changes in genome expression are triggered at the earliest stages of interaction, probably on contact with the penetrating juveniles. However, the conferred protection does not restrict J2 penetration or movement inside the roots; conversely, it somehow restrains the building of feeding sites and the opportunity of J2s to become sedentary and develop. Further investigation is needed to promote the practical use in the field of plant protection by BCAs, because of the complex interactions that such beneficial microorganisms may have with existing soil microbiome and with metabolisms of different plant species. However, biological control of nematodes through plant activation seems a potential suitable low-impact management strategy that can be profitable for farmers, diffused in organic agriculture, and compatible with EU agricultural policy.

## Acknowledgements

The authors want to thank Mr. Ahmed El Bahrawy (IPSP-CNR, Italy) for his technical assistance

## Author Contributions

SM and PL conceived, designed, and performed the experiments, and analyzed the data; SM wrote the paper.

